# GFAP splice variants fine-tune glioma cell invasion and tumour dynamics by modulating migration persistence

**DOI:** 10.1101/2021.08.19.456978

**Authors:** Jessy V. van Asperen, Rebeca Uceda-Castro, Claire Vennin, Jacqueline Sluijs, Emma J. van Bodegraven, Andreia S. Margarido, Pierre A.J. Robe, Jacco van Rheenen, Elly M. Hol

## Abstract

Glioma is the most common form of malignant primary brain tumours in adults. Their highly invasive nature makes the disease incurable to date, emphasizing the importance of better understanding the mechanisms driving glioma invasion. Glial fibrillary acidic protein (GFAP) is an intermediate filament protein that is characteristic for astrocyte- and neural stem cell-derived gliomas. Glioma malignancy is associated with changes in GFAP alternative splicing, as the canonical isoform GFAPα is downregulated in higher-grade tumours, leading to increased dominance of the GFAPδ isoform in the network. In this study, we used intravital imaging and an *ex vivo* brain slice invasion model. We show that the GFAPδ and GFAPα isoforms differentially regulate the tumour dynamics of glioma cells. Depletion of either isoform increases the migratory capacity of glioma cells. Remarkably, GFAPδ-depleted cells migrate randomly through the brain tissue, whereas GFAPα-depleted cells show a directionally persistent invasion into the brain parenchyma. This study shows that distinct compositions of the GFAP-network lead to specific migratory dynamics and behaviours of gliomas.

## Introduction

Glioblastoma multiforme (GBM, grade IV glioma) is the most common and most aggressive tumour of the central nervous system, with an incidence of 3 per 100,000 people and a crude median survival of 9 months after diagnosis^1^. GBM is currently incurable and this is for a large part due to the highly invasive nature of glioma cells^2–4^. Standard-of-care treatment for GBM consists of surgical tumour resection, followed by chemo- and radiotherapy, but fails to fully eradicate highly invasive glioma cells. As a consequence, patients often relapse after treatment and the tumour rapidly re-grows.

The intermediate filament (IF) protein glial fibrillary acid protein (GFAP) is a signature type III IF protein of glioma cells that has been implicated in tumour migration^5–7^. The role of IFs in glioma invasion and migration has only gained attention recently^8^. With over 70 genes encoding different IF proteins, the IF family is one of the largest human gene families and IF expression patterns are highly cell- and tissue type-specific^9^. Changes in the composition of the IF-network are associated with alterations in malignancy. For example, during the epithelial-to-mesenchymal (EMT) transition, a process linked to increased cellular invasiveness and cancer progression^10^, the IF-network of cancer cells with an epithelial origin changes from a keratin-dominant to a vimentin-dominant network^11–13^. In addition, breast cancer invasion is linked to changes in the IF-network, with a switch from keratin 8 to keratin 14 expression in invasive cells^14^. GFAP is an IF protein that is classically used to identify malignancies of glial origin, such as astrocytomas and glioblastomas^15^. In addition to GFAP, gliomas can heterogeneously express a combination of IFs including vimentin, synemin, and nestin^16^, which are located within the same filament in the cell^17^. GFAP is differentially spliced, and GFAPα and GFAPδ are the two isoforms that are most highly expressed and best studied. The GFAPδ isoform results from alternative splicing with a 3’ polyadenylation event, where the last two exons 8 and 9 of GFAPα are replaced by exon 7a, leading to an alternative 42 amino acid C-terminal tail^18,19^. The two isoforms have different assembly properties^20^, protein interactions^19,21^, and differ in their expression patterns, with GFAPα predominantly expressed in mature astrocytes and GFAPδ in the neurogenic niches of the human brain^22,23^.

In previous studies, we and others have shown that glioma malignancy is associated with alterations in GFAP splice isoform levels^6,24–29^. As such, RNA sequencing analysis of the cancer genome atlas (TCGA) database showed that increasing glioma malignancy grades are associated with a lower overall expression of GFAP and a shift towards higher levels of the alternative splice variant GFAPδ relative to GFAPα^6^. Increasing the GFAPδ/α ratio *in vitro* leads to an upregulation of genes encoding proteins that are involved in the interactions between cells and the extracellular matrix (ECM) such as laminins, integrins, and matrix metalloproteinase 2 (MMP-2)^5–7,20^. Besides, immunohistochemical analysis of glioma tissue samples linked GFAPδ expression to an altered cellular morphology^26,28^ and to more invasive tumours based on neuroimaging^27^. Although these observations are suggestive for changed glioma cell behaviour upon alterations in GFAP-isoform expression, a full characterization of changed behaviour has not yet been performed.

In this study, we investigated how manipulation of GFAP-isoform expression affects human glioma cell invasion and growth dynamics *ex vivo* and *in vivo*. We longitudinally monitored the growth patterns of a total of twelve clones of U251-MG glioma cells depleted from either the GFAPα or the GFAPδ isoform in *ex vivo* organotypic mouse brain slices and in mouse brains *in vivo* with intravital imaging. We show that manipulation of the GFAP-network strongly affects the motility of glioma cells and tumour growth patterns. GFAPδ-KO cells form denser tumours, have increased motility compared to control tumours and migrate randomly, whereas GFAPα-KO cells show a more diffuse growth pattern and migrate more persistently towards the brain parenchyma.

## Results

### GFAP-isoform expression differs between low- and high-grade gliomas

Using differential gene expression analysis of RNA sequencing data from the TCGA database, we previously showed that the ratio of splice variants GFAPα and GFAPδ differs between low grade- and high-grade gliomas^6^. Since this publication, 37 additional patient samples were included in the TCGA database. We therefore re-analysed the RNA sequencing data of the updated TCGA cohort and confirmed our previously reported findings. Whereas canonical splice variant GFAPα was significantly decreased in grade IV glioma compared to lower grades glioma (grade II and III) (Supp. Fig. 1a), the expression of alternative splice variant GFAPδ was not different between the different grades (Supp. Fig. 1b). Thus, there is an increased dominance of GFAPδ in high-versus lower-grade glioma, as illustrated by the significant increase in the GFAPδ/α ratio (Fig. 1a).

**Figure 1.**
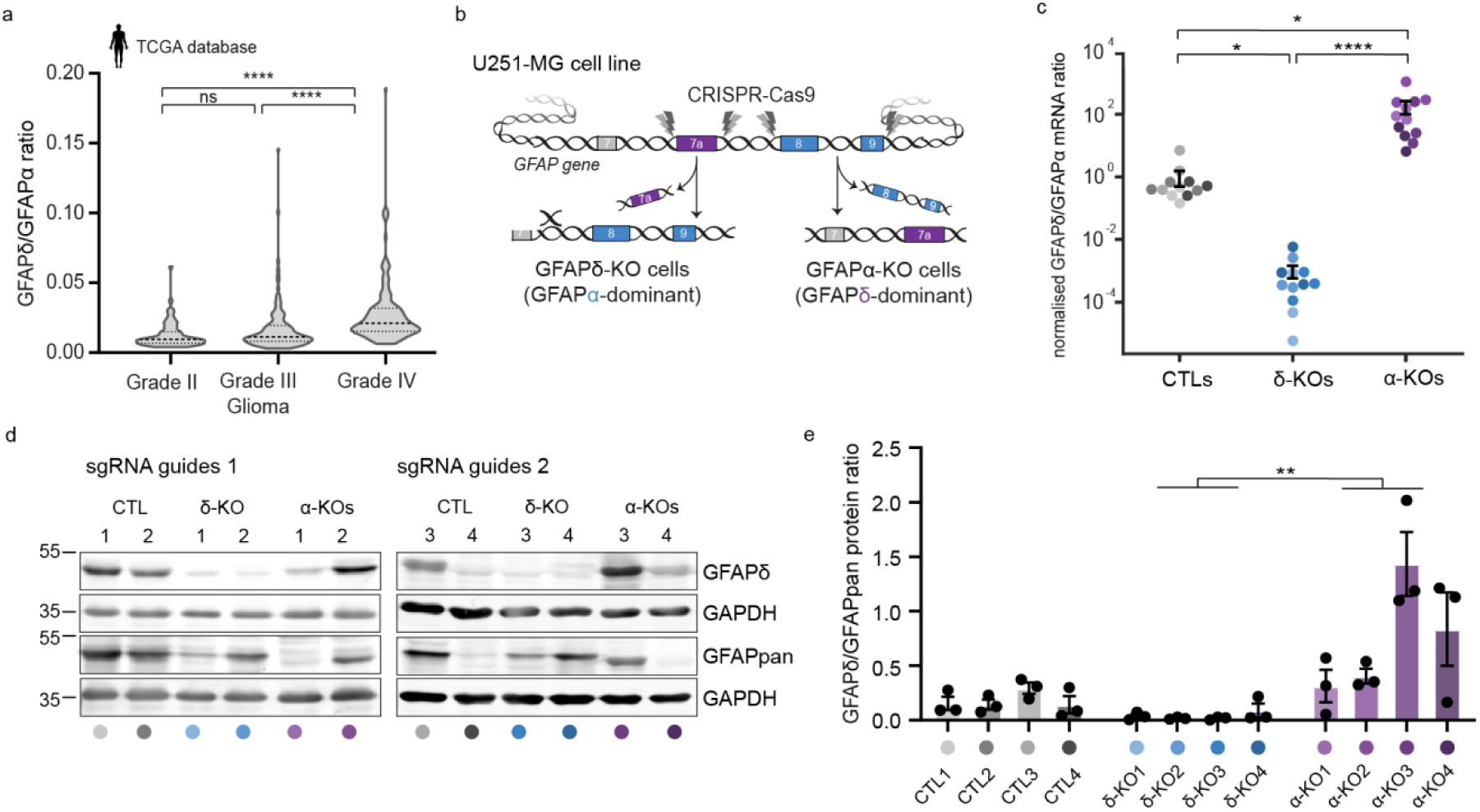
GFAPδ/GFAPα ratio in the TCGA database and generation of GFAP isoform KO lines to regulate the GFAPδ/ GFAPα ratio in U251-MG glioma cells. (a) Violin plots of the GFAPδ/GFAPα ratio in tumour samples of grade II (n= 64), grade III (n= 130), and grade IV (n= 153) astrocytoma, obtained from normalised isoform expression data of the TCGA database. Significance was determined using a Kruskal-Wallis test followed by a Dunn’s multiple comparisons test. (b) Schematic illustration of the GFAP gene with the Cas9-targeted locations to generate GFAPδ- and GFAPα-KO lines. GFAPδ-KO and GFAPα-KO lines were generated using two sets of sgRNAs, and four clones per group were selected and characterised, leading to a total of 12 lines. (c) GFAPδ/GFAPα mRNA ratio of the GFAP isoform KO lines and controls, represented on a log10 scale. Depletion of exon 7a (GFAPδ-KO) leads to a decrease in the GFAPδ/ GFAPα ratio compared to the control cells, whereas depletion of exons 8 and 9 (GFAPα-KO) leads to an increase in the ratio. n=12 biological repeats per group, derived from 4 clones per condition represented with different colour hues. Significance was determined using a Kruskal-Wallis test followed by a Dunn’s multiple comparisons test. (d) Protein levels of GFAPδ and all GFAP isoforms (GFAPpan) in the 12 different lines determined with Western blot. (e) Quantification of GFAPδ/GFAPpan levels in the 12 different lines. Significance was determined using a Kruskal-Wallis test followed by a Dunn’s multiple comparisons test. The data is shown as mean ± S.E.M, *p < 0.05, **p < 0.01, ***p < 0.001, ****p < 0.0001, ns = not significant.

### Modification of GFAP-isoform expression using CRISPR-Cas9

To understand how the different ratios of GFAPδ/α affect the behaviour of the tumour cells, we modified GFAP-isoform expression in the U251-MG human glioma cell line using CRISPR-Cas9 technology, as previously performed in van Bodegraven *et al*., 2019 [ref. 7]. A set of two single guide RNAs (sgRNAs) were used to delete the DNA region encoding the 41 or 42 amino acid tail of GFAPα and GFAPδ, respectively. To create a GFAPα knockout (KO), the intronic regions before and after exon 8 and 9 were targeted, whereas the GFAPδ-KO cells were created by flanking the intronic regions before and after exon 7a (Fig.1b, Supp. Fig. 2b,d). In addition to the previously generated six lines^7^, we engineered six extra clonal lines using a different set of sgRNAs to create a total of four GFAPα-KO lines, four GFAPδ-KO lines, and four control (CTL) lines. Exonal depletion was confirmed with polymerase chain reaction (PCR) (Supp. Fig. 2a,c) and sequencing (Supp. Fig. 2b,d). Targeting the exonic region led to a significant decrease in mRNA levels of the corresponding isoform (Supp. Fig. 2e,f,g) and an increase (GFAPα-KO) or decrease (GFAPδ-KO) of the GFAPδ/α mRNA ratio (Fig. 1c) and the GFAPδ/GFAPpan protein ratio (Fig. 1d,e, Supp. Fig. 2h,i). The clonal lines showed normal IF-network formation, except for GFAPα-KO clone 3, where occasional network collapses were observed (Supp. Fig. 2j). This GFAPα-KO clone 3 had the highest GFAPδ/GFAPpan protein ratio (Fig. 1d), confirming that there is a limit to the level of GFAPδ that can be incorporated into the network^20,30^.

### Depletion of GFAP-isoforms increases cell invasion in *ex vivo* organotypic brain slices

To study how modulation of the GFAP-network affects cell invasion in a physiologically relevant environment, we adapted the *ex vivo* organotypic brain slice model described by Pencheva *et al*., 2017 [ref. 31]. *Ex vivo*, 350 µm thick brain slices of p15-17 mouse pups were prepared and cultured in an air-liquid interface. The twelve clonal lines were transduced with H2B-mNeonGreen to visualise the nuclei and were injected into the lateral ventricles of the organotypic brain slice using a micromanipulator. The H2B-mNeonGreen expressing cells (4 CTLs, 4 GFAPδ-KOs, 4 GFAPα-KOs) were co-injected with an internal control line (I-CTL, CTL clone 1 or 3, Fig. 1d) that expressed H2B-mCherry. The brain slices injected with U-251-MG cells were kept in culture for one week (Fig. 2a). Upon fixation of the *ex vivo* slices, we applied whole-mount immunofluorescent staining for laminin and used RapiClear tissue clearing^32^. Subsequently, we used confocal imaging to create a 3D-reconstruction of the invasion patterns of the cells in the brain slice (Fig. 2b, Supp. Fig. 3). Laminin expression was not only observed along the blood vessels but deposits produced by the glioma cells were also observed at the injection site where the cell density was the highest (Fig. 2b, Supp. Fig. 3). We used this laminin expression pattern to distinguish cells within the tumour core from cells that had invaded into the mouse brain tissue (Supp. Fig. 3).

**Figure 2.**
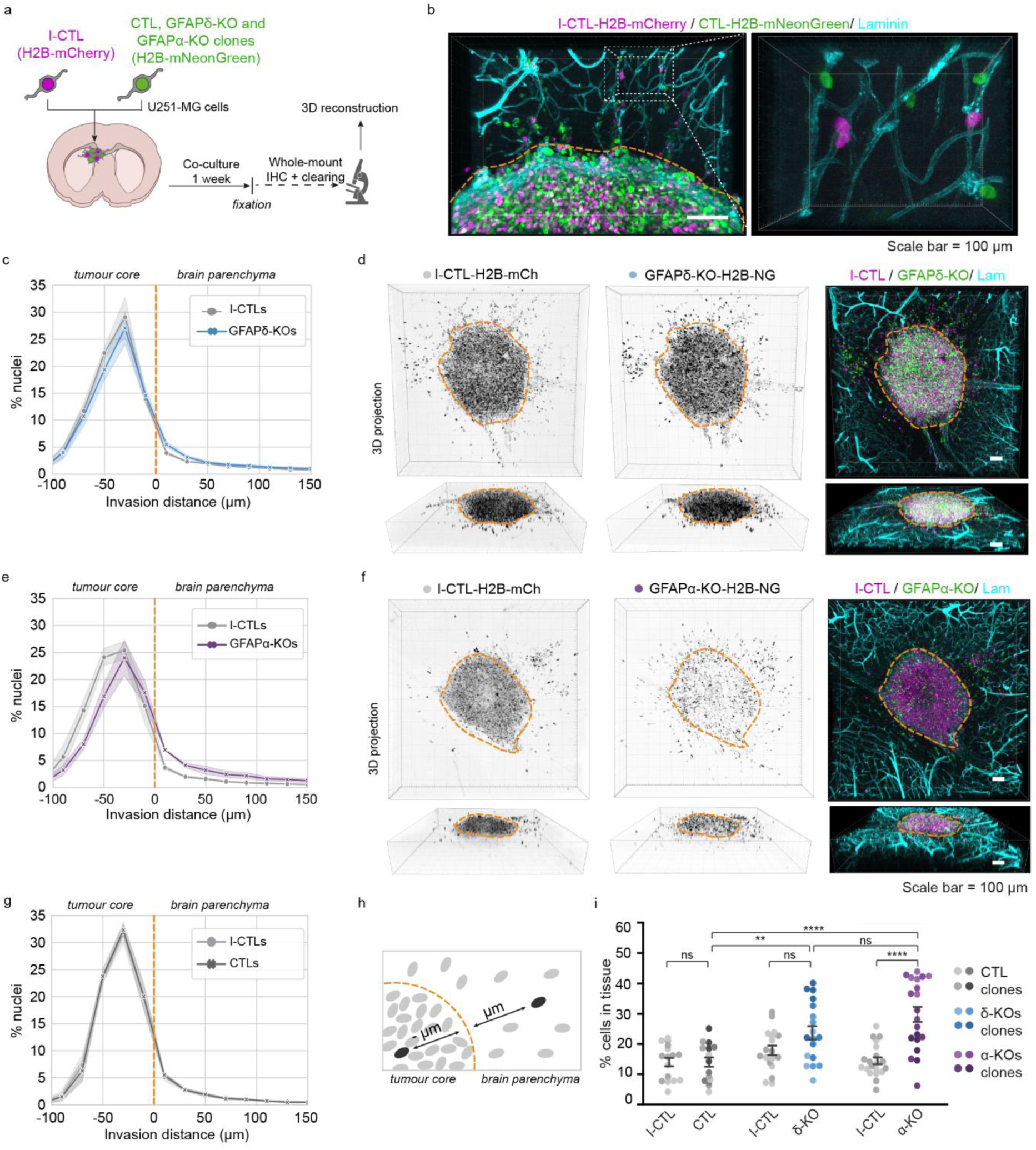
Modification of GFAP isoform expression affects macroscopic growth patterns in organotypic brain slice cultures. (a) Schematic of experimental set-up: H2B-NeonGr expressing control (CTL), GFAPδ-KO and GFAPα-KO clonal lines are injected in organotypic slices together with an H2B-mCherry expressing internal control (I-CTL) and co-cultured for one week. After fixation, whole-mount immunofluorescent staining, clearing and confocal imaging are used to create a 3D reconstruction of the invasion patterns. (b) Representative image of nuclei of an I-CTL (magenta) and a CTL (green) lines within the organotypic slice model. Invading cells are mainly found around the mouse brain vasculature (laminin, cyan). Laminin deposits in the tumour core can be used to distinguish stationary cells from cells invading the tissue, indicated with the orange dotted line. (c) Distribution of nuclei of GFAPδ-KO and I-CTL cell lines in the organotypic slices (n=18 independent experiments, 4 different clones). (d) Representative images of invasion pattern of GFAPδ-KO line 1 and I-CTL 1. (e) Distribution of nuclei of GFAPα-KO cells and I-CTL cells in the organotypic brain slices (n= 20 independent experiments, 4 different clones). (h) Representative image of the invasion pattern of GFAPα-KO line 2 and I-CTL 1. (g) Distribution of I-CTL and CTL nuclei in the organotypic brain slices (n=16 independent experiments, 4 different clones). (h) Schematic depicting the method used to quantify the distribution of nuclei in the organotypic slices. (i) Quantification of the percentage of invaded cells per condition, n= 16 (CTLs), n=18 (GFAPδ-KO), and n= 20 (GFAPα-KO) injected organotypic brain slices derived from 4 different clones per condition. Significance was determined using a two-way ANOVA followed by Tukey’s multiple comparisons test. Scale bar = 100 µm. The data is shown as mean ± S.E.M, *p < 0.05, **p < 0.01, ***p < 0.001, ****p < 0.0001, ns = not significant. NG= mNeonGreen, mCh = mCherry, lam = laminin.

First, we compared the distribution of nuclei of the CTLs, GFAPδ-KOs, and GFAPα-KOs to that of the I-CTLs. We calculated the distance of every individual nucleus from the boundary of the tumour core and plotted the distribution of the cells within different distance bins (Fig. 2h). As expected, the distribution plot of H2B-mNeonGreen CTL lines overlapped with that of the H2B-mCherry expressing I-CTL (Fig. 2g, Supp Fig. 4a). The distribution of nuclei of the GFAPδ-KO cells slightly deviated from the I-CTL line (Fig. 2c,d, Supp Fig. 4b), but the clearest alteration in distribution was observed in the GFAPα-KO cells. Whereas I-CTL cells have the highest density of cells in the tumour core, the GFAPα-KO cells showed a more diffuse growth pattern (Fig. 2f, Supp Fig. 4c). When plotting the distribution of cells, a shift in cell density towards the tumour border and tissue was observed (Fig. 2e), indicating more invasion. We next quantified the percentage of cells in the tissue as a measure for invasion and indeed observed a higher percentage of invading GFAPα-KO cells in comparison to its I-CTL line and in comparison to the CTL lines (Fig. 2i). Whereas GFAPδ-KO had similar percentages of invading cells compared to its I-CTL, a higher percentage of invading cells was measured in comparison to the CTLs (Fig. 2i). This implies that I-CTL cells are more invasive when co-injected with GFAPδ-KO cells, thereby suggesting an indirect effect of the GFAPδ-KO cells on the I-CTL. However, the invasive capacity of the I-CTL lines in the different conditions was not significantly different (Fig. 2i).

To confirm the effect of downregulating GFAPα on tumour distribution patterns, we repeated the *ex vivo* organotypic brain slice invasion experiment with U251-MG cells transduced with an shRNA against the 3’UTR of GFAPα (Supp. Fig. 5a), as earlier published in Moeton *et al.* 2014 [ref. 5]. Targeting GFAPα at the mRNA level led to a diffuse growth pattern and more invading cells, similar to the observations seen in CRISPR-Cas9 modified cells. (Supp. Fig. 5).

### Depletion of GFAPα isoform leads to more diffuse tumours *in vivo*

Next, we aimed to study the GFAP modulated cells in an *in vivo* setting where a functional vasculature is present, and where it is possible to follow tumour progression over time. We used intravital microscopy (IVM), which allows to longitudinally visualise tumour cell behaviour at the single-cell level in a living organism^33^. We separately injected two H2B-mNeonGreen expressing clonal lines with the most extreme GFAPδ/α ratio *per* condition into NOD-Scid IL2Rgnull mice. Tumour development was followed using a cranial imaging window (CIW). An overview image of the tumour at the endpoint was taken, when a well-established tumour had formed (Fig. 3a, b). To quantify the tumour density, we calculated the number of individual cells in the total tumour area. We observed that tumours generated by the GFAPδ-KO were significantly denser than tumours generated by the GFAPα-KO cells (Fig. 3b,c). This suggested that GFAPα-KO cells have a more diffuse growth pattern compared to GFAPδ-KO cells.

**Figure 3.**
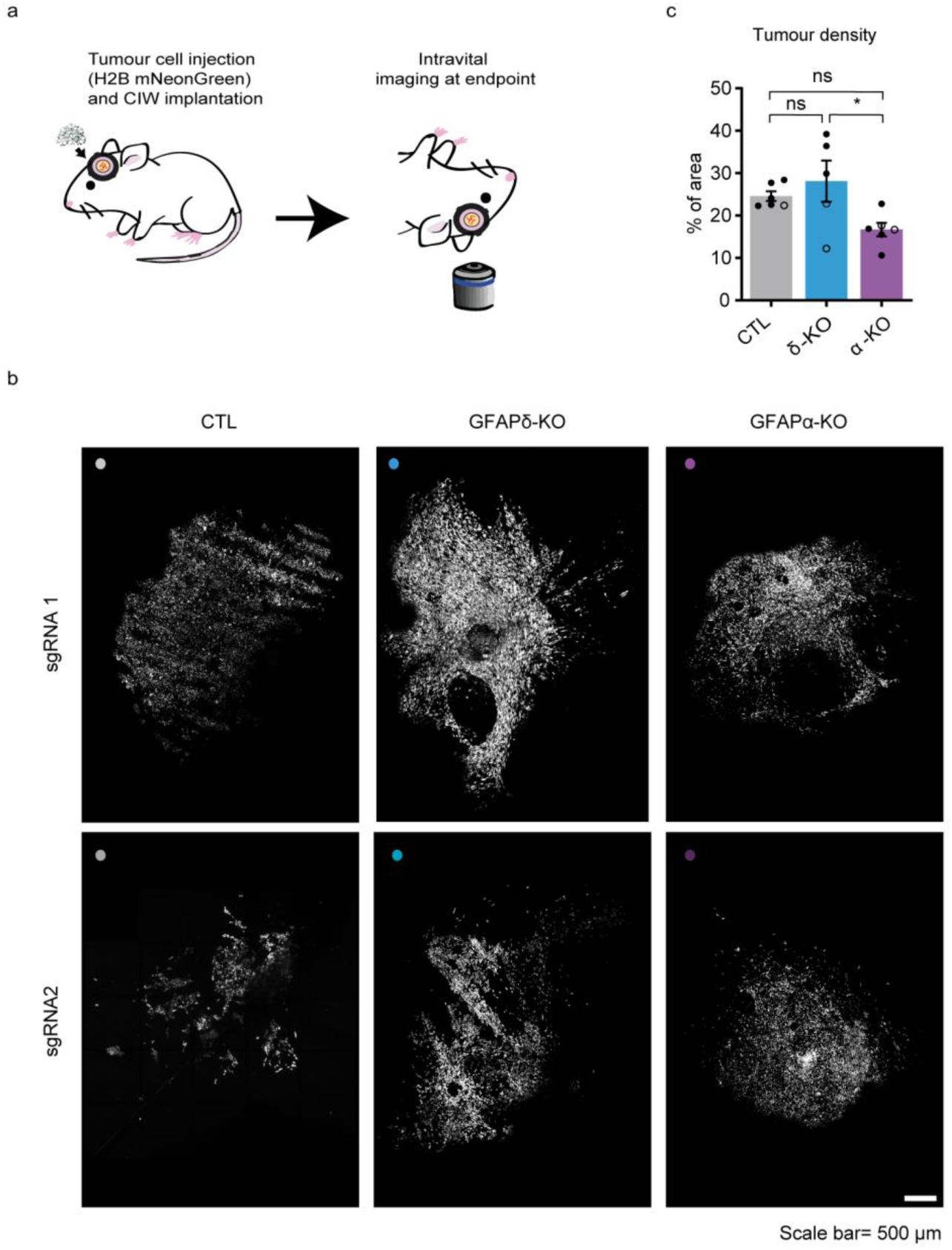
*In vivo* tumour growth dynamics in GFAP modulated tumours. (a) Schematic overview of the experimental setup. U-251-MG GFAP-modulated lines expressing H2B-NeonGreen were implanted in the brain of NSG mice under a CIW. Time-lapse intravital imaging was performed through a CIW to study the tumour growth dynamics of each tumour type. (b) Representative 3D reconstructed tile-scans showing distinct tumours generated by different GFAP-modulated lines. Two clones engineered with different CRISPR-Cas9 guides are presented. Scale bar = 500µm (c) Quantification of tumour density for each indicated tumour type. n=6 (CTLs), n=5 (GFAPδ-KO), and n= 6 (GFAPα-KO) mice. Black dots represent sgRNA1 and white dots represent sgRNA2.The data is shown as mean ± S.E.M, *p < 0.05, **p < 0.01, ***p < 0.001, ****p < 0.0001, ns = not significant, one-way ANOVA followed by Tukey’s multiple comparisons test.

### Depletion of GFAP isoforms increases motility and alters invasion patterns *in vivo*

To further gain insight into the migratory behaviour of GFAP-modulated glioma cells *in vivo*, we again made use of the CIW to longitudinally study invasive behaviours at the single-cell level. At endpoint, a series of time-lapse z-stack images of the tumour was acquired for 6 hours with a time interval of 45 minutes (Fig. 4a). The movement of individual glioma cells was determined by tracking their migration path over time in 3D-reconstructed time-lapse movies (Fig. 4b). Data concerning migration velocity, speed, persistence, and directionality were extracted from the tracks. This showed that depletion of either GFAPδ or GFAPα isoform leads to an increase in the percentage of motile cells compared to the CTL (Fig. 4c). While the GFAPδ-KO cells migrate faster than the CTL cells (Fig. 4d, e), they move with less persistence compared to the GFAPα-KO and CTL cells (Fig. 4f). Considering that directionality is an important factor for invasion, we analysed the directionality patterns in each tumour type and determined whether the cells were migrating towards the tumour core or the brain parenchyma^34^. This demonstrated that GFAPα-KO cells migrate more towards the brain parenchyma while the CTL cells and GFAPδ-KOs migrate more randomly (Fig. 4 b,g). Indeed, this data is in line with our observation that GFAPα-KO tumours are more diffuse than GFAPδ-KO tumours (Fig. 3b, c).

**Figure 4.**
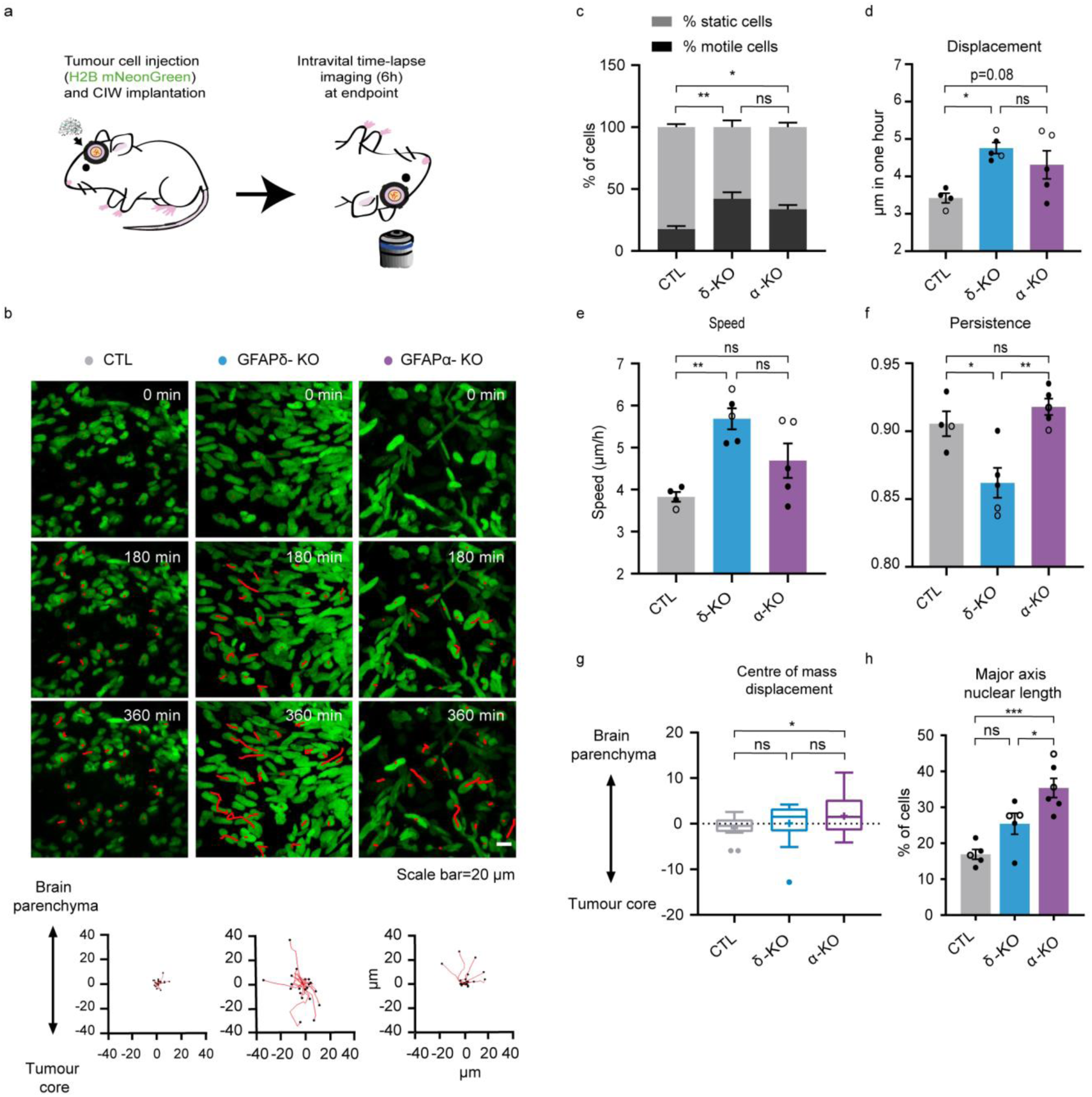
*In vivo* migratory behaviour of tumour cells with different GFAPδ/α ratios. (a) Schematic representation of implantation of CIW and intravital time-lapse imaging over 6 hours with an interval of 45 minutes. (b) Representative still images from a time-lapse movie showing migratory tumour cells in different GFAP modulated tumours. Red lines highlight individual tumour cell tracks. Scale bar=20 µm. Corresponding plots represent tracks from a common origin showing the direction of the tumour cells either towards the tumour core or the brain parenchyma. (c) Percentage of motile (cell displacement > 2 μm/hour) and static cells for each tumour type. (d) Quantification of cell displacement of motile cells for the indicated tumour type. (e) Cell speed of motile cells for the different cell lines (µm/h). (f) Cell persistence of motile cells in the different cell lines. Black dots represent sgRNA1 and white dots sgRNA2. (f) Tukey-style whiskers plot of the centre of mass displacement of individual positions of each condition. (g) Quantification of nuclear cell length in the different cell lines. % of cells with a length higher than 30 µm is represented. n=4 (CTLS), n=5 (GFAPδ-KO), and n=5 (GFAPα-KO) CIW implanted mice from 2 different clonal lines. The data is shown as mean ± S.E.M, *p < 0.05, **p < 0.01, ***p < 0.001, ****p < 0.0001, ns = not significant, one-way ANOVA or two-way ANOVA followed by Tukey’s multiple comparisons test.

It has been recently shown that nucleus stiffness and cell deformability plays an important role in cell motility. For instance, to move in a three-dimensional ECM, the nucleus of a cell must squeeze through the narrow spacing within the brain parenchyma^35–37^. In our model, we observed a significant increase in the nuclear axis length of GFAPα-KOs compared to CTLs (Fig. 4h), which may contribute to the increased ability of GFAPα-KO to infiltrate the brain parenchyma.

## Discussion

The invasive nature of glioma makes the disease highly aggressive and hard to treat. Therefore, precisely understanding the mechanisms driving invasion of glioma cells is crucial for the development of new anti-invasive treatment strategies. In this study, we investigated the role of GFAP-isoforms in glioma cell invasion, using an organotypic brain slice invasion model and intravital imaging through a CIW. We show that the GFAPδ/α ratio affects the macroscopic growth patterns of glioma cells both *ex vivo* and *in vivo* by regulating cell migration speed, directionality, and persistence. Importantly, we demonstrate that GFAPδ-KO cells show increased motility compared to CTL and GFAPα-KO cells *in vivo*, but move randomly. GFAPα-KO cells, on the other hand, move more persistently and have a strong tendency to migrate towards the brain parenchyma. These dynamics of the GFAPα-KO cells lead to a more diffuse infiltration pattern into the brain parenchyma.

Earlier studies that investigated the role of GFAP in cell motility and migration have been somewhat inconsistent. As such, GFAP expression has been linked to both higher^38,39^ and lower^5,40,41^ velocities of cell migration. Also, shRNA mediated knockdown of GFAPα decreased cell velocity in an earlier *in vitro* study^5^, which is inconsistent with the phenotype we describe here. The effect of GFAP depletion on cell behaviour might not only be isoform dependent, but also influenced by the cell-environmental context^34^, which may explain discrepancies between earlier studies performed in 2D. Our study is the first, to our knowledge, to investigate the role of GFAP and its isoforms within the physiological context of the brain. We find that depletion of both isoforms leads to an increase in the number of motile cells, as well as an increase in cell displacement and speed (Fig. 4c,d,e). Therefore, cell-intrinsic motility and velocity may not be dependent on the GFAPδ/α ratio, but on modification of the GFAP-network in general. In contrast to motility and velocity, we demonstrate that directionality and persistence of cell migration is GFAP-isoform dependent. Previous studies have shown that the IF-network can promote migration persistence by modulating microtubule organization and cell polarity^42–44^. Whether the absence of GFAPα or dominance of GFAPδ regulates directional migration similarly remains to be elucidated. The directionality of migration and invasion can also be steered by extrinsic factors^45^ such as ECM composition and topology^45^. For instance, adhesive interaction of the cell with the extracellular microenvironment as well as remodelling of the ECM are required to migrate efficiently through the extracellular space^46^. MMPs are responsible for the degradation of a large range of ECM proteins and GBM cells have been shown to overexpress MMP2 and 9 [ref. 47]. In line with this, we showed in our previous work that modulation of the GFAPα isoform affects genes involved in the compositions of the ECM and extracellular space^6^. In addition, we previously demonstrated that GFAPα-KO cells produce more laminin and overexpress MMP2 [ref. 5,7], and this might contribute to the higher ability of these cells to invade the brain parenchyma persistently.

The findings of this study contribute to our understanding on how a switch in the GFAPδ/α ratio in grade IV glioma patients may affect the aggressiveness of these tumours. Increased dominance of GFAPδ in grade IV glioma tumours has been reported by multiple studies^6,25,26,28^ (reviewed in van Bodegraven *et al.,* 2019 [ref. 29], and was confirmed by analysis of the updated TCGA database (Fig. 1a). Similarly, Brehar and colleagues reported that patients with highly invasive tumours, based on pre-operative MRI, had increased percentages of GFAPδ positive cells^27^. Glioma tumours are known to be highly heterogeneous. This heterogeneity appears to be not only between patients but also between single cells within a tumour^48–50^, therefore it is likely that the same tumour is composed of a mix of cells with a high and low GFAP-δ/α ratio and these distinct cell populations may contribute to different behaviour. Together, it can be hypothesised that a larger population of high GFAPδ/α ratio cells in grade IV tumours contributes to infiltration of the brain parenchyma and subsequent relapse after therapy.

How the shift in GFAP-isoform expression is established in grade IV tumours is currently unknown. Alternative splice events are known to occur more frequently in tumour tissue in comparison to non-malignant cells^51^, and dysregulation of the splicing machinery drives glioma aggressiveness^52^. Recently, it was discovered that hypoxia can induce adult-to-foetal splicing transitions in glioma, regulated by muscle blind-like proteins (MBNL)^53^. Hypoxia is considered an important driver of glioma invasion and is typically associated with grade IV gliomas^54,55^. GFAP has multiple predicted binding motifs for the hypoxic-associated splicing factor MBNL^56^, The link between hypoxia, GFAP alternative splicing, and cell invasion remains to be investigated.

In summary, our work demonstrates the importance of GFAP-isoforms in fine-tuning glioma invasion and tumour dynamics. Together, the increased understanding of the mechanisms driving the invasive behaviours of different GFAP positive populations that form glioma tumours will help develop better anti-invasive therapeutic strategies in the future.

## Materials and Methods

### Cell lines and culture

The cell identity of malignant glioma cell line U251-MG (obtained from Lars Ruether, Institut für Neuropathologie, Universitätsklinikum Münster, Münster, Germany) was confirmed by short terminal repeat analysis (Eurofins Scientifc, Luxembourg city, Luxembourg). All lines were cultured in DMEM high glucose (Gibco 41966052) mixed 1:1 with Ham’s F10 nutrient mix (Gibco 22390025) supplemented with 10% fetal bovine serum (Gibco 10270106/ Biowest S181H) and 1% penicillin/streptomycin (Gibco 15140122) at 37 °C in a humidified incubator with 5% CO_2_. Cell lines were routinely tested negative for mycoplasma contamination.

### Mice

For the generation of organotypic slice cultures, 15-17 day-old C57BL6J mice were used. C57BL6J mice were obtained from Charles Rivers Laboratories and bred in-house. The animals were kept under a normal 12:12h light-dark cycle with lights off at 19:00, at room temperature (21 +/- 2 °C) and at 40-70% humidity conditions, and were fed with chow and water ad libitum.

For intravital imaging experiments, NOD-Scid IL2Rgnull mice (NSG) mice, aged 8 to 20 weeks at the time of cranial window implantation were used. Mice were housed in individually ventilated cage and received food and water *ad libitum*.

All experimental protocols used in this manuscript were in accordance with national regulations and ethical guidelines and were approved by the Centrale Commissie Dierproeven (CCD) and the Instantie voor Dierenwelzijn (IvD).

### TCGA RNA sequencing data collection and analysis

Expression data of GFAP splice variants from the Cancer Genome Atlas (TCGA) was extracted using the TSVdb webtool (http://tsvdb.com)^57^. Normalised RSEM (RNA-Seq by Expectation aximization) count estimates from TCGA Lower Grade Glioma (TCGA-LGG) and glioblastoma multiforme (TCGA-GBMs) projects were extracted and matched with the sample ID to clinical data on histological subtype and malignancy grade downloaded from the TCGA database: https://www.cancer.gov/tcga. The GFAPα and GFAPδ normalised expression levels and GFAPδ/α ratios were compared in data from 64 grade II astrocytomas, 130 grade III astrocytomas and 153 GBMs.

### Generation of CRISPR-Cas9 plasmids

Single guideRNAs (sgRNAs) targeting the intronic regions before and after exon7a (GFAPδ-KO) or exon 8 and 9 (GFAPα-KO) were designed using web resources of the Broad Institute (http://tools.genome-engineering.org/)^58^ or CRISPOR.org (http://crispor.tefor.net/)^59^ and were selected based on proximity to exons and MIT and CFD specificity score^60^. The sgRNA complementary oligonucleotide templates were cloned into pSpCas9(BB)-2A-Puro (Addgene, #48139) or pSpCas9(BB)-2A-GFP (Addgene, #48138) plasmids after BbsI (Thermo Fisher Scientific) digestion. Plasmids were isolated using a Maxiprep kit (LabNed) and the sequence was verified by Sanger sequencing (Macrogen, Amsterdam, The Netherlands). Per GFAP-isoform, two pairs of sgRNA plasmids were generated (Table S1). The sgRNA pairs cloned into the pSpCas9(BB)-2A-Puro plasmid (sgRNAs 1) and the clonal cell lines generated with these plasmid pairs have been described in van Bodegraven et al., 2019 [ref. 7]. The sgRNA pairs cloned into the pSpCas9(BB)-2A-GFP plasmid (sgRNAs 2) are first described in this paper [ref. 7].

For cell transfection of the CRISPR-Cas9 construct and clonal expansion, U251-MG cells were seeded at a density of 0.8 to 1.2 x 10^5^ cells in an uncoated 6-well plate. Twenty-four hours after seeding, two CRISPR-Cas9 plasmids (1 μg DNA total) containing sgRNA upstream and downstream of the targeted exons of the GFAP isoforms were co-transfected using polyethylenimine (PEI, 166 ng/mL final concentration). Cells transfected with the pSpCas9(BB)-2A-Puro plasmid (sgRNAs 1) were treated with 1 μg/mL puromycin (Sigma-Aldrich, 58-58-2) 24 hours after transfection and were selected for 96 hours. The drug-resistant pool was expanded and clonal lines were generated by single-cell sorting cells into 96-well plates using fluorescence-activated cell sorting (FACS; FACSAria II Cell Sorter). Cells transfected with the pSpCas9(BB)-2A-GFP plasmid (sgRNAs 2) were selected for GFP using FACS (FACSAria II Cell Sorter) 48 hours after transfection. The GFP-positive pool was expanded and clonal lines were generated by plating cells at low densities in 96-well plates (0.5 cell/well). The 96-well plates were inspected for colony formation and clonal lines were expanded. Control lines were generated by selecting clonal lines of cells transfected with the empty and undigested plasmids.

### Selection of CRISPR-Cas9 targeted clonal lines

PCR screening was used to identify clonal lines in which the targeted DNA region in the GFAP gene was depleted. Genomic DNA was isolated from cell pellets of the clonal lines. Cells were lysed in 5 mM Tris HCl (pH 8.8) at 95°C for 10 minutes and treated with proteinase K at 56°C for 30 minutes. The CRISPR-Cas9 targeted DNA region was amplified using primers described in Table S1, using the FirePol PCR Master Mix (Solis BioDyne, 04-12-00S15). PCR products were separated on a 1.5% agarose gel containing SYBR Safe (Thermo Fisher Scientific, S33102) and GFAP-isoform KO lines were identified based on the presence of predicted smaller PCR products. Depletion of the targeted DNA region was confirmed by isolating the amplified DNA of the PCR product using the PureLink Quick Gel Extraction Kit (Thermo Fisher Scientific, K210012) and Sanger sequencing (Macrogen, Amsterdam, The Netherlands).

### shRNA construct design

Lentiviral shRNA expression plasmids targeting GFAPα or non-targeting controls were generated as described by Moeton et al., 2014 [ref. 5]. In short, lentiviral shRNA expression plasmids from the RNAi Consortium (TRC) Mission library were obtained from Sigma-Aldrich (TRCN0000083733)^61^. A human shRNA construct against nucleotides 2674-2694 in the 3’ untranslated region of the GFAPα transcript or a SHC002 non-targeting construct (NTC) with no homology to human sequences were cloned into the pLKO.1 backbone.

### Lentiviral production and transduction of cells

Lentiviruses encoding NTC or GFAPα shRNA were produced as described by Moeton et al., 2014 [ref 5]. U251-MG cells were transduced with lentiviral particles encoding NTC or GFAPα shRNA with a multiplicity of infection (MOI) of 0.5. Three days after transduction, cells were selected by treatment with 1 μg/mL puromycin (Gibco, A1113803) to create stable cell lines.

All U251-MG GFAP-modulated cell lines (with CRISPR-Cas9 or shRNAs) and controls were transduced with lentiviruses to induce expression of H2B-mNeonGreen or H2B-mCherry. The pLV-H2B-mNeonGreen-IRES-puro plasmid was a gift from Dr. Hugo Snippert^62^, the pLenti6-H2B-mCherry plasmid was a gift from Torsten Wittmann (Addgene plasmid # 89766). Lentiviral particles were produced with standard third-generation lentiviral protocol. In short, 2 x 10^7^ 293T cells (ATCC, ATCC-CRL-11268) were plated in a 15 cm^2^ dish and transfected the next day with a total of 51.6 μg DNA of an envelope plasmid (pMD2.G), packaging plasmids (pMDLg/pRRE and pRSV-Rev) and pLV-H2B-mNeonGreen-IRES-puro or pLenti6-H2B-mCherry plasmid using PEI (166 ng/mL final concentration). The medium was replaced 24 hours after transfection. After 48 hours, the medium containing virus particles were collected and filtered through a 0.22 μm filter. The supernatants were ultracentrifuged at 22,000 rpm (rotor 70Ti, Beckman ultracentrifuge) at 16 °C for 2 hours and 40 minutes. The pellet was resuspended in PBS + 0.5% BSA (Sigma), aliquoted and stored at −80°C until further use. The viral titre was determined by transducing 293T cells with a dilution series of the virus. The viral titre was estimated in transducing units (TU) / mL by counting the number of transduced fluorescent cells 48 hours after transduction. The GFAP modulated cells were transduced with H2B-mNeonGreen and H2B-mCherry lentiviral particles with an MOI of 1. Cells were passaged once and positive cells were selected by keeping the cells in medium containing 1.5 µg/mL puromycin (H2B-mNeonGreen clones) or 10 µg/mL blasticidin (H2B-mCherry clones) for three days.

### Western blot analysis

Total protein was extracted from cultured cells scraped in suspension buffer [0.1M NaCl, 0.01 M Tris HCl (pH 7.6), 0.001 M EDTA, and Complete EDTA-free protease inhibitor cocktail (Roche)] and sonicated (2x 10 seconds) in an ultrasonic bath. An equal amount of 2x SDS loading buffer [100 µM Tris (pH 6.8), 4% SDS, 20% glycerol, 5% 2-ME, and bromophenol blue] was added to the cell suspension, samples were heated at 95 °C for 5 minutes and DNA was broken down by pushing the sample through a 25-gauge needle. Equal amounts of sample were loaded on a 10% SDS-page gel and proteins were separated by electrophoresis. Proteins were then blotted on a 0.45-µm pore size nitrocellulose membrane (GE Healthcare) using a wet/tank transfer blotting system (Biorad, 170390). Membranes were blocked in blocking buffer (50 mM Tris pH 7.4, 150 mM NaCl, 0.25% (w/v) gelatin, 0.5% Triton-X100) for 10 minutes and incubated with primary antibodies (Table S2) in blocking buffer overnight at 4 °C. Membranes were washed with TBS with 1% Tween (TBS-T) three times for 10 minutes and then incubated with secondary antibodies (Table S2) in blocking buffer at room temperature for 1 hour. After three washing steps with TBS-T and one washing step with MilliQ, the membrane blots were scanned with the Odyssey Clx Western Blot Detection System (Li-Cor Biosciences). The background-corrected signal intensity of bands corresponding to the GFAPpan and GFAPδ proteins were measured and normalised against the intensity levels of glyceraldehyde 3-phosphatedehydrogenase (GAPDH) bands on the same blots.

### RNA isolation, cDNA isolation and real-time quantitative PCR

For RNA extraction of cultured cells, cells were seeded on poly-D lysine (PDL)-coated wells of a 24-well plate at a density of 4 × 10^4^ cells. After three days in culture, cells were lysed in TRIzol (Thermo Fisher Scientific, 15596026) and RNA was extracted using standard TRIzol-chloroform extraction methods. RNA concentration and purity were measured using Varioscan Flash (Thermo Fisher Scientific). 200 to 500 ng of RNA were used to prepare cDNA using the QuantiTect Reverse Transcription Kit (Qiagen, 205311) according to the manufacturer’s protocol. The generated cDNA was used for real-time quantitative PCR using the SYBR Green Master mix in a QuantStudio 6 Flex Real-Time PCR system (Thermo Fisher Scientific, 4309155) using the primers listed in Table S1. Expression values were calculated by transforming Ct values (2^-Ct^) and were normalised to the mean value of the transformed Ct values of the reference genes GAPDH and Alu element Jurka (Alu- J).

### Immunocytochemistry

For immunocytochemistry on cultured cells, cells were seeded on PDL-coated coverslips in a 24-well plate at a density of 2 × 10^4^ cells. After three days in culture, the cells were fixed in 4% paraformaldehyde (PFA) dissolved in phosphate buffer saline (PBS), pH 7.4 for 30 minutes. Cells were washed in PBS, incubated in a blocking buffer (50 mM Tris pH 7.4, 150 mM NaCl, 0.25% (w/v) gelatine, and 0.5% triton X-100) at room temperature for 15 min, and afterwards with primary antibodies (Table S2) in blocking buffer overnight at 4°C. Coverslips were washed with PBS and incubated with secondary antibodies (Table S2) and Hoechst 33528 (1:1000, Thermo Fisher Scientific, H3569) in blocking buffer at room temperature for 1 hour. After washing steps with PBS, the coverslips were mounted on microscopy slides with Mowiol (0.1 M tris-HCl pH 8.5, 25% glycerol, 10% Mowiol (Merck Millipore, 81381). The samples were imaged using a Zeiss Axioscope A1 microscope with a 40x objective.

### Generation of Organotypic Brain Slices

For the generation of organotypic brain slices, the protocol of Pencheva et al., 2017 was adapted^31^. Postnatal day 15 – 17 C57BL6J pups were decapitated, the brains were dissected and captured in ice-cold artificial cerebrospinal fluid (aCSF, pH7.2: 10 mM Hepes, 21 mM NaHCO3, 1.2 mM NaH_2_PO_4_, 2.5 mM KCl, 2 mM MgCl_2_, 2 mM CaCl_2_, 5 mM D-glucose, 250 mM glycerol in milliQ). The brains were transferred to a petri dish and cerebellum and olfactory bulbs were removed. The cerebrum was glued to the vibratome cutting stage using a drop of Loctite 401 glue (Henkel Adhesives) with the rostral part facing upwards. The vibratome cutting stage was mounted on a VT1000S vibratome (Leica Biosystems,1404723512) and tissue was fully submerged in carbonated ice-cold aCSF. Coronal brain slices of 350 μm were cut with a speed of 0.1 mm/s and a frequency of 7 Hz. Slices with visible lateral ventricles were transferred to 1.0-μm porous membrane inserts (Corning®, 353102) in a 6-well plate with slicing medium [DMEM:F12 (Gibco, 11320), 1% L-Glutamine (Gibco, 25030123), 5 mM HEPES, 21 mM NaHCO_3_ and 1% pen/strep (Gibco, 15140122], with a maximum of 4 slices per transwell insert. Residual aCSF was removed from the inserts, the brain slices were washed with PBS and the transwells were transferred to 1.5 mL recovery medium (DMEM:F12, 25% FBS, 1% L-Glutamine, 5 mM HEPES, 21 mM NaHCO_3_ and 1% P/S) below the transwells, allowing the slices to be cultured at the air-liquid interface. The slices were cultured at 37 °C in a humidified incubator with 5% CO_2_ overnight. The next day, the transwells were dipped twice in PBS and transferred to a 6-well plate containing NSC medium (DMEM:F12 – glutamax, 1% pen/strep, 10 ng/mL EGF (Peprotech, AF-100-15-A), 10 ng/mL FGF (Peptrotech, AF-100-18B) before injection of cells.

### Organotypic brain slice invasion assay

Cells were counted using the Countess 3 FL Automated Cell Counter (Thermo Fischer Scientific, AMQAF2000) and suspensions of 2.5 × 10^5^ cells/ μl were prepared. For the mixed cell injections, H2B-mNeonGreen expressing cells were mixed at a 1:1 ratio with H2B-mCherry expressing internal control cells. For the CRISPR-Cas9 modulated cells, CTL-1-H2B-mCherry was used as an internal control (I-CTL1) for the sgRNAs 1 lines (CTL 1 and 2, GFAPδ-KO 1 and 2, GFAPα-KO 1 and 2), and CTL-3-H2B-mCherry was used as an internal control (I-CTL2) for the sgRNAs 2 lines (CTL 3 and 4, GFAPδ-KO 3 and 4, GFAPα-KO 3 and 4). For the shRNA modulated cells, NTC-H2B-mCherry was used as an internal control. A Hamilton 0.5 μL syringe model 7000.5 KH (Hamilton, 86250) was assembled on a Narishige micromanipulator model MM-3 (Narishige group) and was placed on the magnetic board of a Leica MS5 dissection microscope (Leica Biosystems), using a Narishige GJ-8 magnetic stand (Narishige group). The syringe was rinsed with acetone, 70% ethanol, and PBS before use. Before injection, the cell suspension was mixed and 0.5 μL was taken up by the syringe. The needle was placed into the lateral ventricle of the brain slice by moving 50 μm into the tissue and 40 μm out. The cell suspension was slowly injected into the lateral ventricle of the mouse brain tissue, filling up the lateral ventricle without overflowing on the tissue. The medium of the organotypic brain slices was replaced every 2-3 days. One week after injection of the cells, the brain slices were washed with PBS and fixed in 4% PFA in PBS at 4 °C overnight.

### Whole-mount immunofluorescence and clearing

For immunohistochemistry staining of the whole organotypic brain slice, the 3DISCO immunostaining protocol was adapted^63^. The porous membrane surrounding the brain tissue was cut out and transferred to a 24-well dish. The tissue was permeabilised in 2% Triton-X-100 (Roche, 40319421) in PBS and subsequently incubated in PBSGT blocking solution (0.2% gelatin, 0.5% Triton-X-100, 0.01 % thimerosal or 0.2 % sodium azide in 1x PBS), for a minimum of 4 hours at room temperature. Primary antibodies were diluted in PBSGT + 0.1% saponin, 300 μL was added to the tissue and incubated on a horizontal shaker (70 rpm) at 37 °C for 3 days. Tissue injected with mixed H2B-mNeonGreen/ H2B-mCherry cells were incubated with rabbit anti-laminin antibodies (1:1000, Table S2). The empty wells were filled with PBS to avoid evaporation of the primary antibody mix. The tissue was washed 6 times in PBS with 0.25% Triton-X-100 for one hour. Secondary antibodies were diluted in PBSGT + 0.1% saponin and the mix was spun down to precipitate aggregates. 300 μL was added to the tissue and incubated on a horizontal shaker (70 rpm) at 37 °C for 24h. The tissue was washed 6 times in PBS with 0.25% Triton-X-100 for one hour. For tissue clearing, the brain slices were transferred to iSpacers (SunJin Lab Co., #IS002) mounted on microscope slides and 300 μL of RapiClear 1.47 (SunJin Lab Co, #RC147001) was added on the brain slices. The slices were cleared at 37 °C on a horizontal shaker (30 rpm) for 45 minutes, mounted with a coverslip in RapiClear 1.47, and sealed with transparent nail polish. The cleared brain slices were imaged using an LSM 880 (Zeiss) confocal microscope equipped with a 3-channel QUASAR Detection Unit (000000-2078-293). The entire population of injected cells was imaged with a 10x objective (N-Achroplan 10x, 420940-990-000) at 1.77 μm pixel resolution Z-plane increments of 6.63 μm and using image tiling. Smaller regions were imaged using a 20x objective (LD Plan-NEOFLUAR 20x, 421350-9970-000) at 0.42 μm pixel resolution and 3.39 μm Z-plane increments.

### Quantification of cell invasion in *ex vivo* slices

Cell invasion in the organotypic brain slices was quantified in the confocal generated images using ImageJ (1.53c) and Imaris software (version 8.4). Upon blinding, images were excluded from analysis when errors had occurred during the injections of cells (overflowing of tissue, large populations of unhealthy looking cells). Using ImageJ software, image tiling was used to reconstruct the entire population of injected cells. The tiled z-stack consisted of an H2B-mNeonGreen and H2B-mCherry channel with the nuclei of the injected glioma cells and a laminin channel staining the mouse vasculature. In addition to staining the vasculature, laminin also gave rise to a diffuse staining at the location where tumour density was highest, as shown by H2B-mCherry/ H2B-mNeonGreen signal. This staining of the ECM deposits generated by the tumour cells was used to draw a boundary between the tumour core and the mouse tissue (Supp. Fig.3) in the different z-planes. An additional channel was generated in which only the tumour core laminin staining was selected. The stitched images with additional laminin-channel were imported into the Imaris Software (version 8.4). The ‘create surface’ function was used to generate a 3D surface of the laminin tumour core signal (background subtraction, estimated diameter 17.8, threshold =2, voxels =1). The ‘create spots’ function was used to generate individual spots (11 μm + PSF-elongation along the Z-axis) of the H2B-mCherry nuclei using a standardised Quality threshold filter. Using the same function, the same number of H2B-mNeonGreen spots was generated. The ‘distance transformation’ function was used for the tumour core surface, generating a new channel where the intensity of the signal represented the distance from the ‘outside surface object’ or ‘inside surface object’. The Imaris software was used to calculate for every H2B-mCherry and H2B-mNeonGreen nucleus the distance to the tumour core, using the ‘intensity center’ calculation within the “statistics” function. The excel file was exported and histograms of the distances (bin size 20 μm) were created using the NumPy package of the Python software^64^.

### Tumour cell injection and cranial window implantation (CWI) surgery

Two clones per condition generated by two different sgRNA were used for the *in vivo* experiments: CTL 1 and 3, GFAPδ-KO 2 and 3, and GFAPα-KO 2 and 4. Clones with the most extreme GFAPδ/α ratio were selected, except for the GFAPα-KO clone 3 as network collapses were observed in this line. Per injection, 100,000 U-251-MG cells were resuspended in 3μl of PBS and injected the same day as the cranial window was implanted. CWI was performed as previously described^65^. In short, mice were sedated with 4% isoflurane inhalation for inducing anaesthesia and 1.5-2% during surgery. The hair from the back of the neck up to the eyes was shaved. Next, the mouse head was firmly fixed with ear bars in a stereotaxic device. Eye ointment was applied to prevent the animal’s eyes from drying out. Next, the skin was cut circularly. After scraping the periosteum underneath to the edges of the skull, a circular groove of 5 mm diameter was drilled over the right parietal bone. After craniotomy, the dura mater was removed with a fine forceps. Next, tumour cells were injected stereotactically using a 10 μl Hamilton syringe with a 2 pt style needle in the middle of the craniotomy at a depth of 0.5 mm. The exposed brain was sealed with silicone oil and a 6 mm coverslip glued on top. Dental acrylic cement (Vertex) was applied on the skull surface to cover the edge of the coverslip and a 3D printed plastic ring was glued around the coverslip to provide fixation to the microscope. A single dose of 100 μg/kg of buprenorphine (Temgesic, Indivior Europe Limited) was administered before the surgery and the day after surgery. In addition Rimadyl in water was administered 24 hours before CIW implantation and for a total of 72 hours (Zoetis). After surgery, the mice were provided food and water *ad libitum*. Mice were closely monitored twice per week.

### Intravital imaging

Mice were anaesthetised in an induction chamber with 4.0% isoflurane. Next, they were placed face-up in a custom-designed imaging box. A 3D printed imaging plate facilitated CWI fixation. Isoflurane was introduced through the facemask and ventilated by an outlet on the other side of the box. To study cell migration, time-lapse images of several positions of the tumour volume were acquired every 45 minutes for a maximum of 6 hours, during which the climate chamber surrounding the microscope was kept at 37 °C and the mouse body temperature was monitored with a rectal thermometer. For each position, images of the complete z stack of the tumour were acquired, with a step size of 3 µm. Imaging was performed on an inverted Leica SP8 multiphoton microscope with a chameleon Vision-S (Coherent Inc., Santa Clare, CA, www.coherent.com). This microscope is equipped with a 25x (HCX IRAPO NA0.95 WD 2.5mm) water objective with four non-descanned detectors (NDDs). The NDDs detected the following wavelengths: NDD1 <455 nm, NDD2 455–505 nm, NDD3 500–550 nm, NDD4 555–680 nm. H2B-mNeonGreen was excited with 944 nm and detected with NDD3. Scanning was performed in a bidirectional mode at 400 Hz and 12 bit, with a zoom of 1, and 512 × 512 pixels.

### Tracking migration of tumour cells

The tracking of migratory cells was done as previously described^65^. After imaging, acquired z-stacks were corrected for z and xy shifts with Huygens Professional software program (version20.10). Up to 300 cells per mouse were tracked manually with an ImageJ plugin (“MTrackJ” Rasband, W.S., ImageJ, U. S. NIH, Bethesda, Maryland, USA). At the start of each movie, a random cell was selected. The XY position was determined over time and the displacement, speed and persistence for each cell were calculated by Excel (Microsoft).

The spatial average of all cell positions was used to measure the centre of mass displacement. For each border position, the centre of mass along the Y-axis was measured by the ‘Chemotaxis and Migration Tool’. Calculation of the centre of mass (M_end_). i= index of single cells, n= number of cells, X_i,end_ Y_i,end_ = coordinates of the respective endpoint.

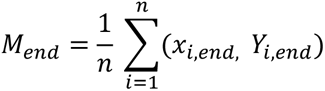

### Statistical analysis

The normality of data was tested using the Shapiro-Wilk test. For all normally distributed measurements, one-way ANOVA (when >2 means were compared) or two-way ANOVA followed by Tukey’s multiple comparisons test were used to determine significance, set to p < 0.05. For non-normally-distributed measurements, a Kruskal-Wallis test (when >2 means were compared) followed by Dunn’s multiple comparisons test were used to determine significance. All p values were two-tailed. Levels of significance were set as follows: ns > 0.05, *0.05 ≤ p > 0.01, **0.01 ≤ p > 0.001, ***0.001 ≤ p > 0.0001, ****p ≤ 0.0001. Error bars are presented as mean ± S.E.M. All statistical analyses were performed using GraphPad Prism software (version 9.1.2, GraphPad Software, USA).

## Acknowledgements

The results shown here are partly based on data generated by The Cancer Genome Atlas Research Network (https://www.cancer.gov/tcga). The authors thank all members of the Hol and van Rheenen labs for thoughtful discussion. This study was funded by the Dutch Cancer Society [KWF 101123, J.v.A, R.U.C, J.v.R., E.H], a HFSP fellowship (C.V.), the Portuguese Foundation for Science and Technology (FCT, GABBA program-PD/BD/105748/2014, A.S.M.), the T and P Bohnenn Foundation (P.R) and the Josef Steiner Foundation (J.v.R).

## Author Contributions

J.v.A. and R.U.C. performed conceptualization, data collection, formal analysis, methodology, and wrote the original draft. C.V. performed conceptualization, methodology, supervision, and reviewed and edited the manuscript. J.S. performed data collection, methodology, and reviewed and edited the manuscript. E.v.B. performed conceptualization, methodology, and reviewed and edited the manuscript. A.S.M. performed methodology, and reviewed and edited the manuscript. P.R. performed conceptualization, methodology, supervision, and reviewed and edited the manuscript. J.v.R. performed conceptualization, funding acquisition, project administration, methodology, supervision, and reviewed and edited the manuscript. E.H. performed conceptualization, funding acquisition, project administration, methodology, supervision, and reviewed and edited the manuscript.

## Data availability statement

All data generated and analysed in this study are included in the manuscript and its supplementary files.

## Supplementary Figures

**Supp. Fig. 1.**
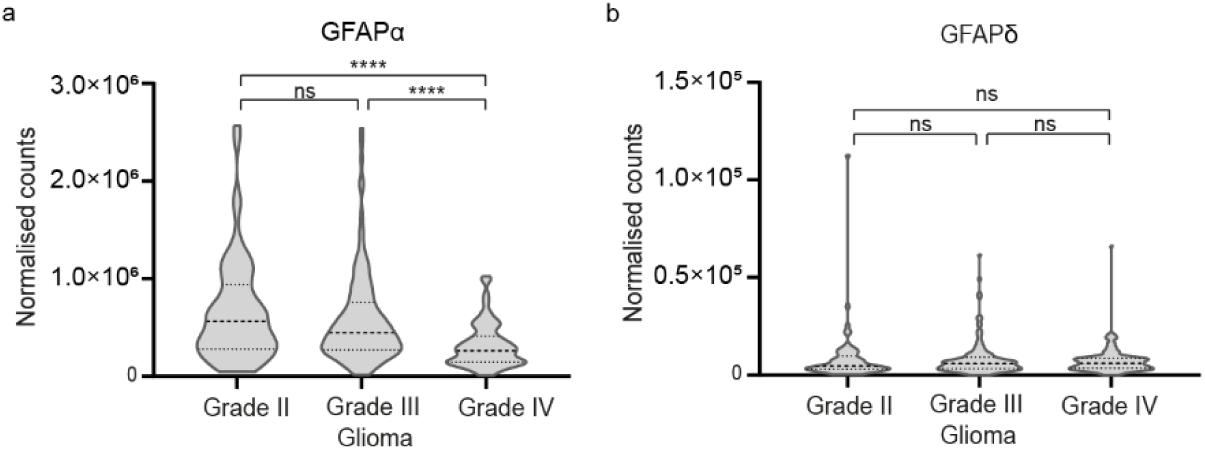
GFAP isoform expression in different grades of astrocytoma. Violin plots of GFAPα and GFAPδ levels in tumour samples of grade II (n= 64), grade III (n= 130), and grade IV (n= 153) astrocytoma, obtained from normalised isoform expression data of the TCGA database. GFAPα levels (a) are decreased in grade IV astrocytoma, whereas GFAPδ levels (b) remain constant. Significance was determined using a Kruskal-Wallis test followed by Dunn’s multiple comparisons test. The data is shown as mean ± S.E.M, *p < 0.05, **p < 0.01, ***p < 0.001, ****p < 0.0001, ns = not significant.

**Supp. Fig. 2.**
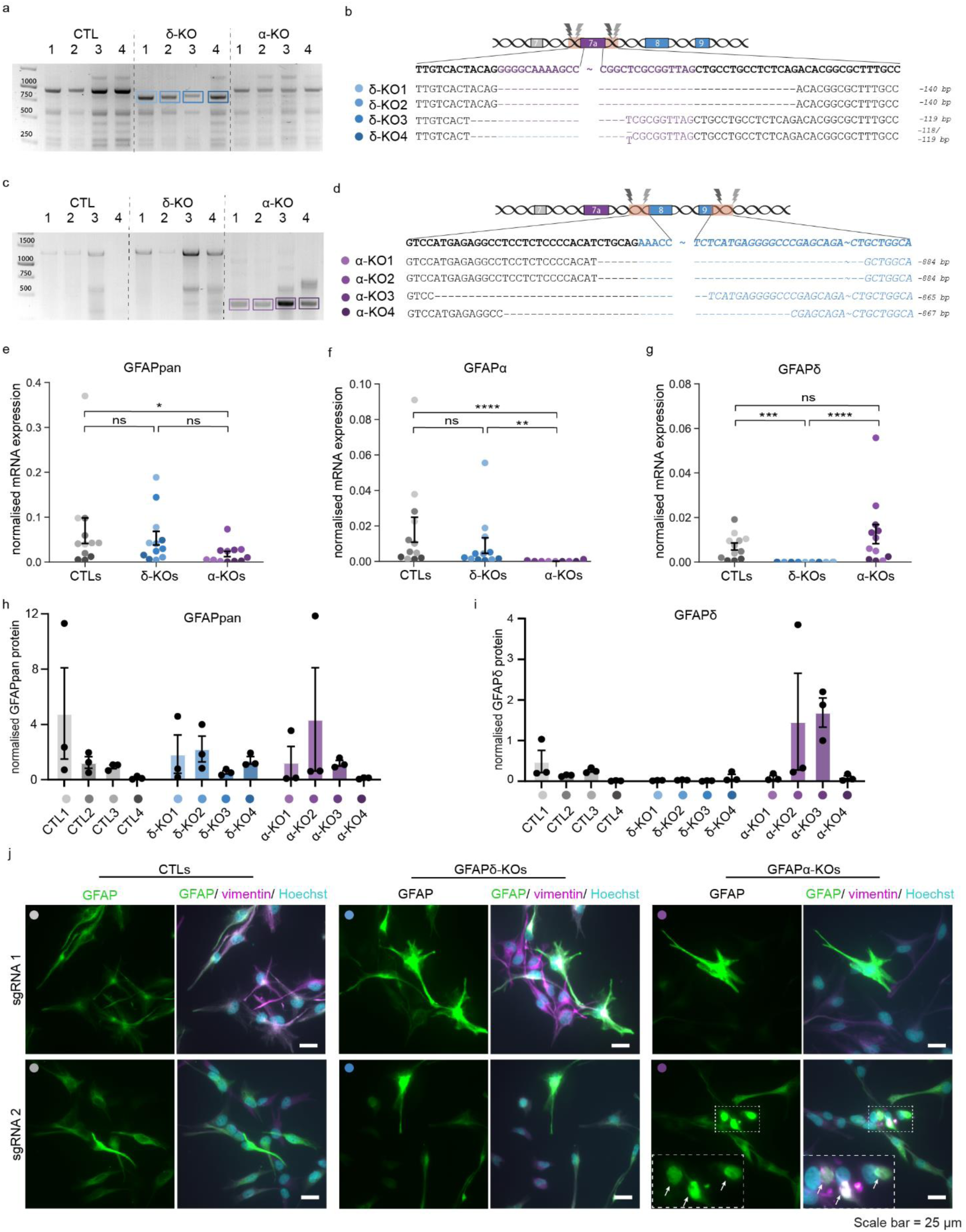
Characterisation of the GFAP modulated cells. (a,b) PCR amplification and sequencing of the GFAP gene around exon 7a. GFAPδ-KO lines have a deletion of 140 bp (GFAPδ-KO 1 and 2, sgRNAs 1) or 118/119 bp (GFAPδ-KO 3 and 4, sgRNAs 2) of exon 7a. (c,d) PCR amplification and sequencing of the GFAP gene around exons 8 and 9. GFAPα - KO lines have 884 bp deletion (GFAPα-KO 1 and 2, sgRNAs 1) or 865/867 bp deletion (GFAPα-KO 3 and 4, sgRNAs 2) of exons 8 and 9. (e-g) mRNA levels of GFAPpan (e), GFAPα (f), and GFAPδ (h) normalised against GAPDH and AluJ. Deletion of exon 7a (GFAPδ-KO lines) leads to a significant reduction in GFAPδ mRNA levels, but not to a reduction in GFAPα or GFAPpan. Deletion of exons 8 and 9 (GFAPα-KO lines) leads to a significant reduction in both GFAPα and GFAPpan levels, but not in GFAPδ mRNA levels. n= 12 individual experiments per group, derived from 4 clones per condition represented with different colour hues. Significance was determined using a Kruskal-Wallis test followed by Dunn’s multiple comparisons test. (h,i) Protein levels of GFAPpan (g) and GFAPδ (h) normalised against GAPDH. (j) The GFAP-network in six different clonal lines (CTL-1, CTL-3, GFAPδ-KO 2, GFAPδ-KO 3, GFAPα-KO 2, GFAPα-KO 3) shown with immunofluorescence. GFAP is integrated in the IF-network (shown with vimentin) in all lines. IF-network collapses were occasionally observed in GFAPα-KO line 3, indicated with white arrows. Scale bar = 25 μm. The data is shown as mean ± S.E.M, *p < 0.05, **p < 0.01, ***p < 0.001, ****p < 0.0001, ns = not significant.

**Supp. Fig. 3.**
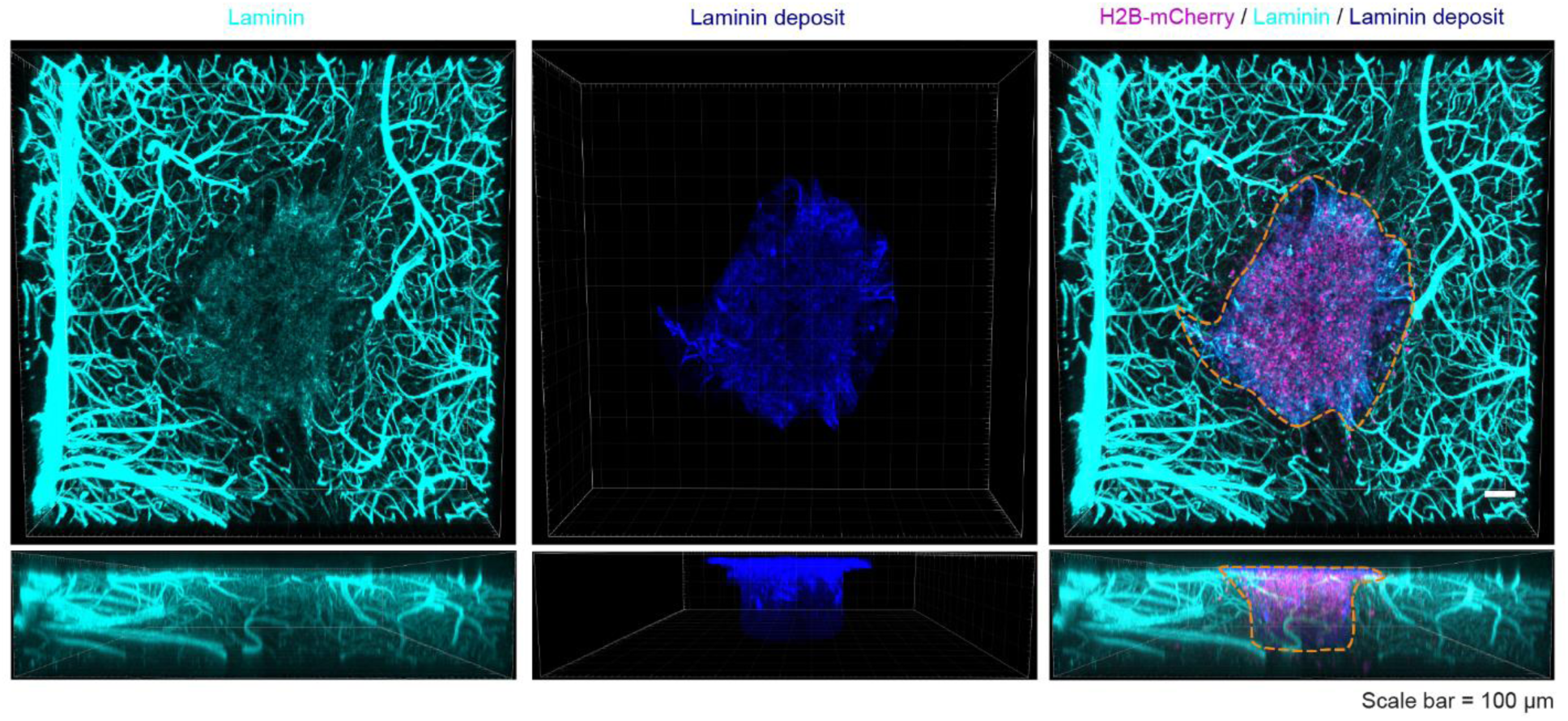
Laminin staining can be used to distinguish tumour core from invading cells. Overexposure of laminin reveals background staining that colocalizes with the highest density of H2B-mCherry nuclei at the site of injection. This laminin deposit is used to distinguish cells in the tumour core *versus* cells that invaded the tissue, indicated with the orange dotted line. Scale bar = 100 μm.

**Supp. Fig. 4.**
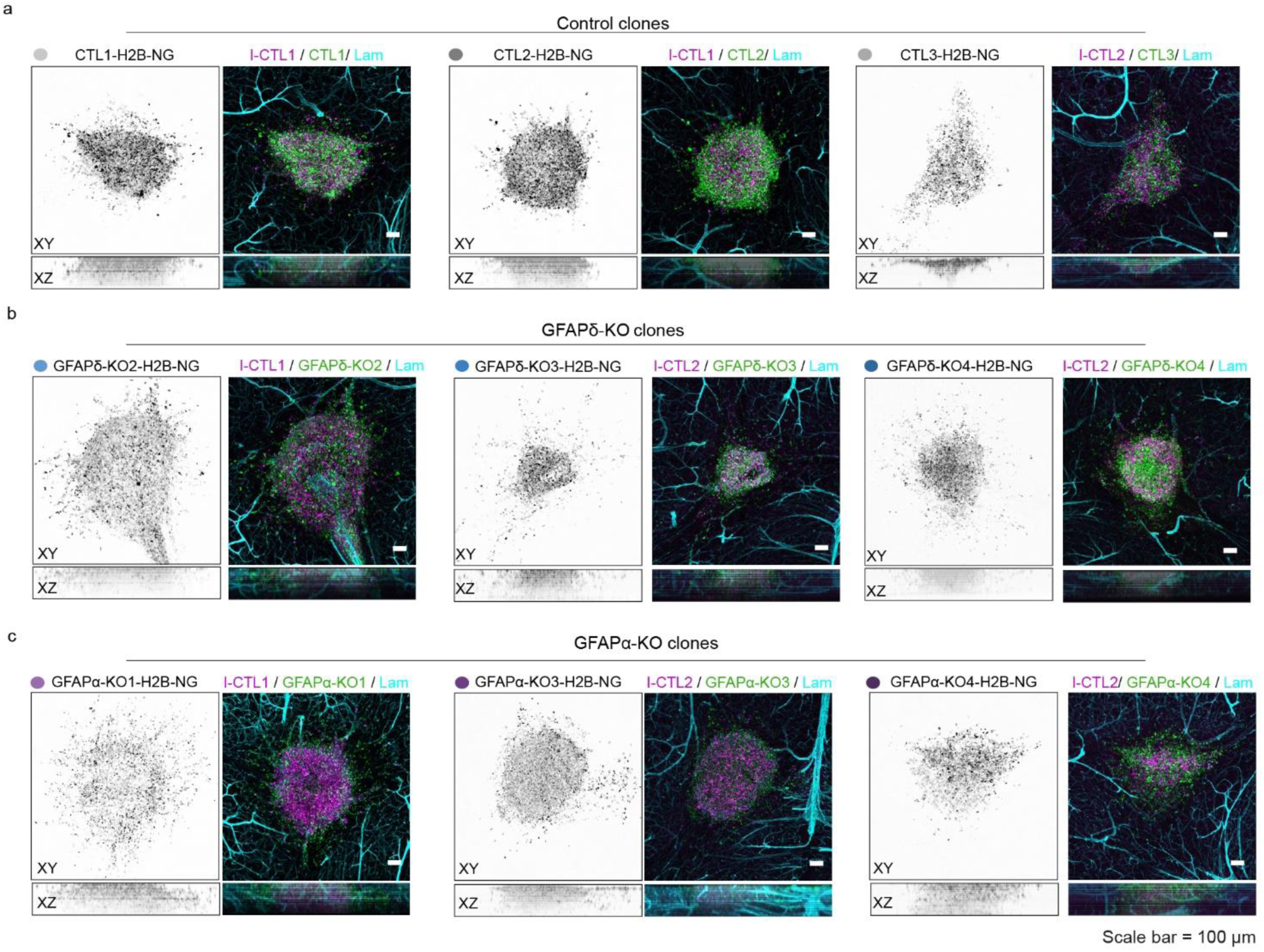
Representative images of organotypic slice cultures injected with the different clonal lines. Representative images of injected I-CTL-H2B-mCherry cells together with H2B-mNeonGreen expressing CTL clones (a), GFAPδ-KO clones (b) or GFAPα-KO clones (c). Scale bar = 100 μm. NG= mNeonGreen, Lam = laminin.

**Supp. Fig. 5.**
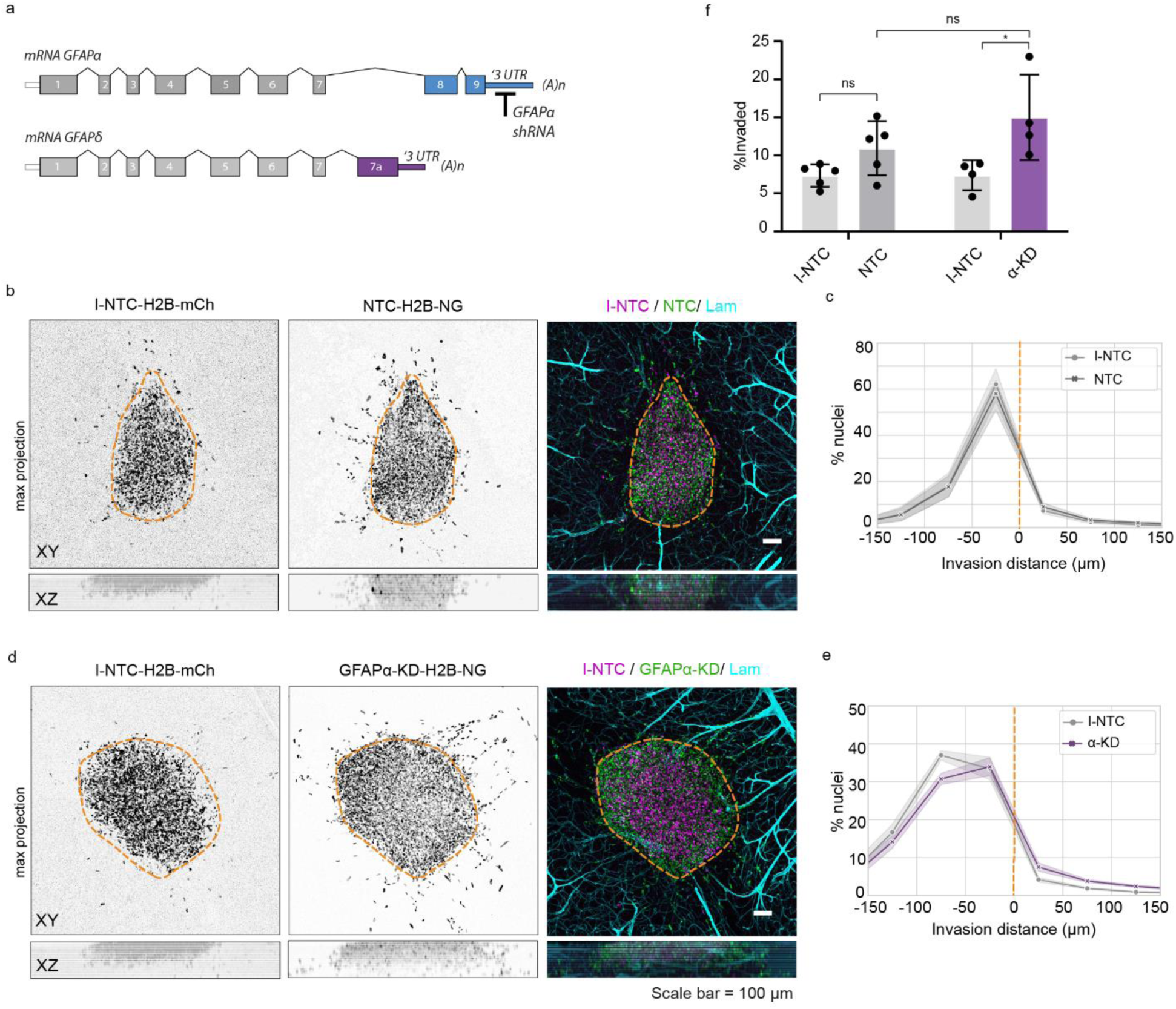
Invasion patterns of GFAPα-KD cells. (a) Schematic illustration of GFAPα shRNA target site. (b,c) I-NTC and NTC show similar distribution patterns of nuclei in organotypic brain slices. Histograms show the percentage of nuclei per 50 μm bins, with negative values representing cells within the tumour core and positive values representing cells in the mouse brain tissue (n= 5 independent experiments). (d,e) GFAPα-KD cells show a more diffuse growth pattern in comparison to I-NTC cells and a shift in cell distribution towards the mouse tissue (n=4 independent experiments). (f) Quantification of the percentage of invaded cells. GFAPα-KD cells show higher percentages of cell invasion in comparison to the I-NTCs, but not in comparison to the NTCs. Significance was determined using a two-way ANOVA followed by Tukey’s multiple comparisons test. Scale bar = 100 μm. The data is shown as mean ± S.E.M, **p < 0.05, **p < 0.01, ***p < 0.001, ****p < 0.0001, ns = not significant. I-NTC = internal non-targeting control, NTC = non-targeting control, NG = mNeonGreen, mCh = mCherry, Lam = laminin, UTR = untranslated region.

## Supplementary Tables

**Table S1.**
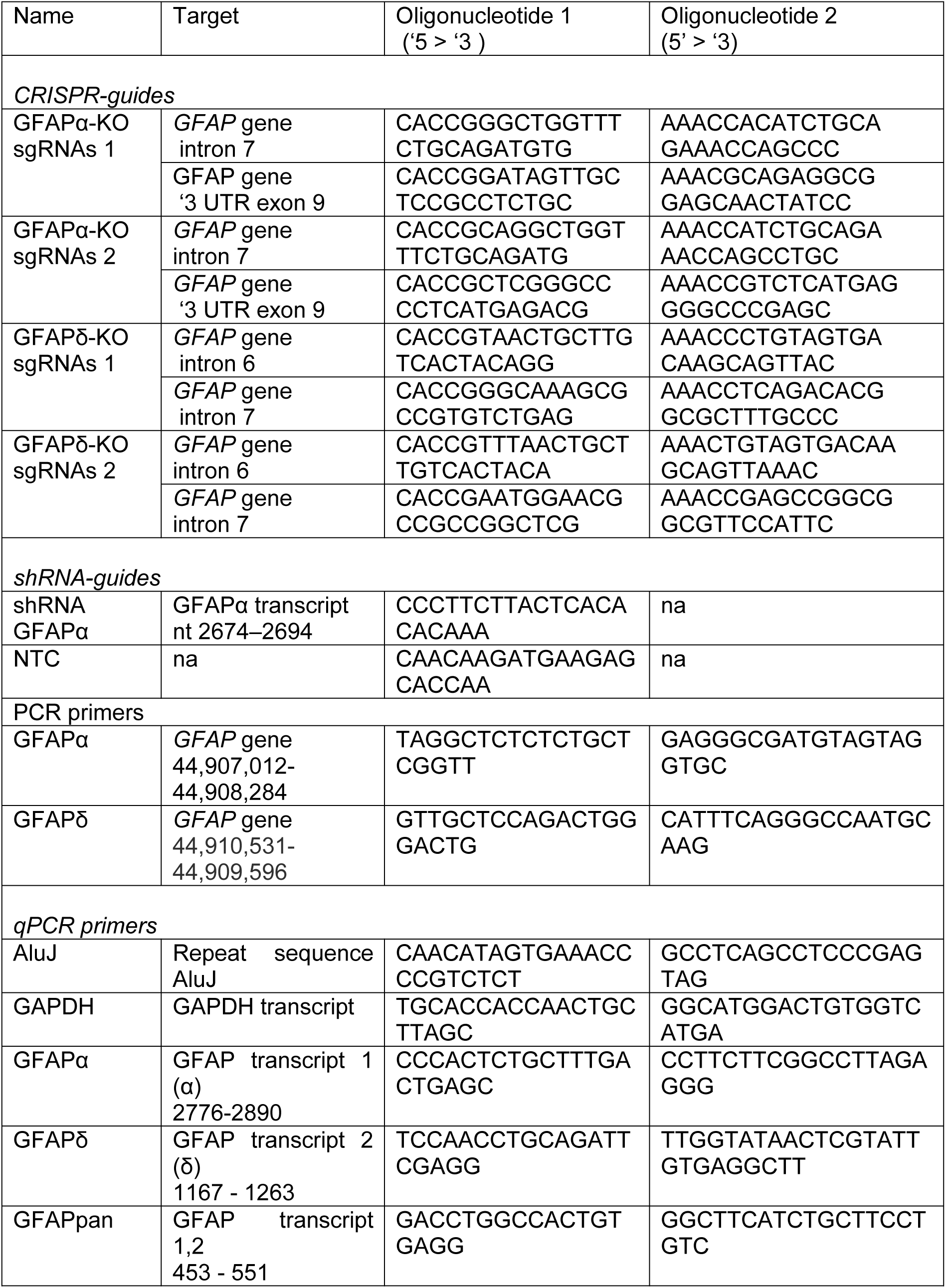
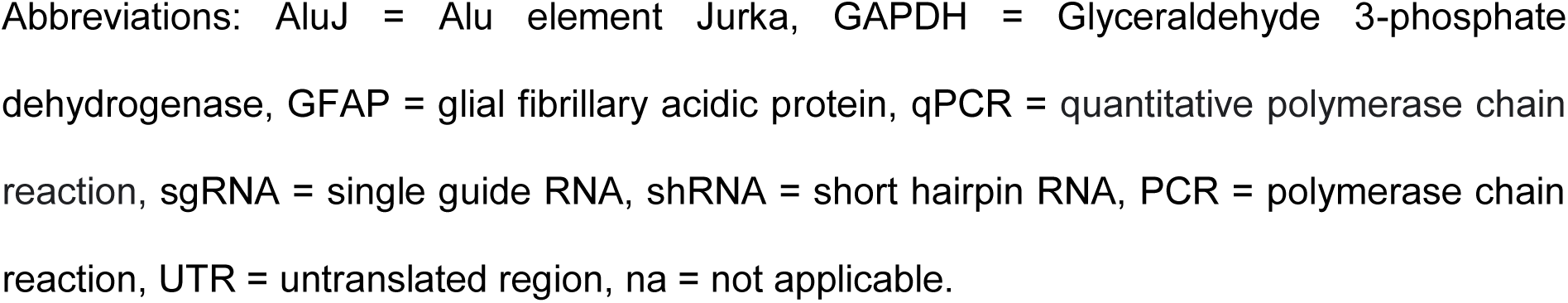
Overview of oligonucleotides.

**Table S2.** List of antibodies.

| Antibody | Product number, Company | (WM)-IF | WB |
| --- | --- | --- | --- |
| Chicken anti-vimentin | AB5733, Chemicon, Temecula, CA, USA | 1:1500 | - |
| Mouse anti-GAPDH | MAB374, Chemicon, Temecula, CA, USA | - | 1:2000 |
| Rabbit anti-GFAP | #Z0334, Dako (Agilent), Santa Clara, CA, USA | 1:1000 | 1:50000 |
| Rabbit anti-GFAPδ | Manufactured in house. Bleeding date: 27-11-2003 (purified in 2004) | - | 1:2000 |
| Rabbit anti-laminin | L9393, Sigma Aldrich, St Louis, MO, USA | 1:1000 | - |
| Donkey anti-chicken Cy3 | 703-175-155, Jackson Immuno Research, West Grove, PA, USA | 1:1000 | - |
| Donkey anti-mouse AF647 | 703-606-150, Jackson Immuno Research, West Grove, PA, USA | - | 1:2000 |
| Donkey anti-rabbit Cy3 | 711-166-152, Jackson Immuno Research, West Grove, PA, USA | 1:1000 | - |
| Donkey anti-rabbit AF647 | 711-606-152, Jackson Immuno Research, West Grove, PA, USA | 1:1000 | - |
| Goat anti-rabbit IRDye800 | 926-32211, LI-COR, 4647 Superior Street Lincoln, NE, USA | - | 1:5000 |

Abbreviations: AF= Alexa Fluor, GAPDH= Glyceraldehyde 3-phosphate dehydrogenase, GFAP = glial fibrillary acidic protein, WB = Western blot, (WM)-IF = (whole-mount)-immunofluorescence

